# Autistic traits, but not schizotypy, predict overweighting of sensory information in Bayesian visual integration

**DOI:** 10.1101/230003

**Authors:** Povilas Karvelis, Aaron R. Seitz, Stephen M. Lawrie, Peggy Seriès

## Abstract

Recent theories propose that schizophrenia/schizotypy and autistic spectrum disorder are related to impairments in Bayesian inference i.e. how the brain integrates sensory information (likelihoods) with prior knowledge. However existing accounts fail to clarify: i) how proposed theories differ in accounts of ASD vs. schizophrenia and ii) whether the impairments result from weaker priors or enhanced likelihoods. Here, we directly address these issues by characterizing how 91 healthy participants, scored for autistic and schizotypal traits, implicitly learned and combined priors with sensory information. This was accomplished through a visual statistical learning paradigm designed to quantitatively assess variations in individuals’ likelihoods and priors. The acquisition of the priors was found to be intact along both traits spectra. However, autistic traits were associated with more veridical perception and weaker influence of expectations. Bayesian modeling revealed that this was due not to weaker prior expectations but to more precise sensory representations.

## Introduction

In recent years Bayesian inference has come to be regarded as a general principle of brain function that underlies not only perception and motor execution, but hierarchically extends all the way to higher cognitive phenomena, such as belief formation and social cognition. Impairments of Bayesian inference have been proposed to underlie deficits observed in mental illness, particularly schizophrenia^1–3^ and autistic spectrum disorder (ASD)^4–7^. The general hypothesis for both disorders is that the weight, also called “precision”, ascribed to sensory evidence and prior expectations is imbalanced, resulting in sensory evidence having relatively too much influence on perception.

In schizophrenia, overweighting of sensory information could explain the decreased susceptibility to perceptual illusions ^8^, as well as the peculiar tendency to jump to conclusions ^9^. Moreover, the systematically weakened low-level prior expectations might lead to forming compensatory strong and idiosyncratic high-level priors (beliefs), which would explain the emergence and persistence of delusions as well as reoccurring hallucinations ^1–3^.

In ASD, the relatively stronger influence of sensory information could explain hypersensitivity to sensory stimuli and extreme attention to details. The weaker influence of prior expectations would also result in more variability in sensory experiences. The desire for sameness and rigid behaviors could then be understood as an attempt to introduce more predictability in one’s environment ^4^. Furthermore, this could lead to prior expectations which are too specific and which do not generalize across situations ^5^. While all theories agree that the relative influence of prior expectations is weaker in ASD, the primary source of this imbalance is debated: does it arise from increased sensory precision (i.e. sharper likelihood) or from reduced precision of prior expectations? ^10–12^ (**Fig. 1**). Some authors argue for attenuated priors^4, 11^, while others argue for increased sensory precision ^6, 7, 10, 13^ but conclusive experimental evidence is lacking.

A number of studies have aimed at testing Bayesian theories, either in a clinical population, or by studying individual differences in the general population^14–17^ under the hypothesis of a continuum between autistic/schizotypal traits and ASD/schizophrenia ^18–20^. Attenuated slow-speed priors were reported in a motion perception task in individuals with ASD traits ^14^. Autistic children also showed attenuated central tendency prior in temporal interval reproduction^21^.

**Figure 1.**
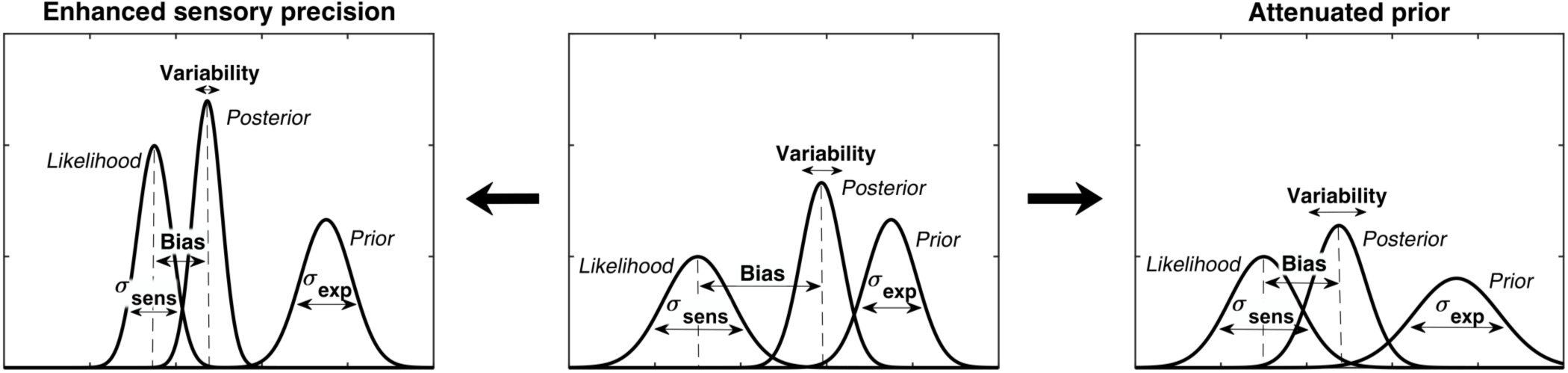
Alternative hypotheses for ASD impairments within the Bayesian inference framework. In Bayesian terms, the percept can be described as a posterior distribution, which is a combination of sensory information (likelihood) and prior expectations (prior). Two contrasting hypotheses have been proposed to underlie behavioural differences in ASD: enhanced sensory precision, i.e. smaller σ_sens_ (left) vs. attenuated priors, i.e. larger σ_exp_ (right). Both hypotheses predict a reduced influence (bias) of the prior on the location of the posterior distribution (posterior mean). However, these alternatives differ in their predictions for perceptual variability (posterior width): the enhanced sensory precision hypothesis should lead to reduced variability while the attenuated prior hypothesis should lead to increased variability. By measuring both bias and variability, our experimental paradigm can distinguish between these two hypotheses.

Attenuated priors were also reported in perceptual tasks that incorporate probabilistic reasoning ^15, 22^. However, the direction of gaze priors ^23^ and the light-from-above priors ^24^ were found to be intact. Autistic children also demonstrated intact ability to update their priors in a volatile environment in a decision-making task ^25^ but a follow-up study in ASD adults showed that they overestimate volatility in a changing environment ^26^.

In schizophrenia/schizotypal traits, Teufel et al.^16^ reported increased influence of prior expectations when disambiguating two-tone images, while Schmack et al.^27,28^ reported weakened influence of stabilizing predictions when observing a bistable rotating sphere.

Overall, the existing findings are not only mixed, but also employ very different paradigms, which makes their direct comparison difficult. Further, a critical limitation of most studies (except for Karaminis et al. ^21^) is the lack of formal computational models that can test whether behavioral differences originate from different priors or from different likelihoods. Moreover, to our knowledge, despite the similarity of the Bayesian theories proposed for ASD and schizophrenia, there is no previous work investigating both autistic and schizotypal traits within the same experimental paradigm so as to test their differences.

We here address these questions empirically in a context of visual motion perception. We used a previously developed statistical learning task^29^ in which participants have to estimate the direction of motion of coherently moving clouds of dots (**Fig. 2**). Chalk et al. ^29^ found that in this task healthy participants rapidly and implicitly develop prior expectations for the most frequently presented motion directions. This in turn alters their perception of motion on low contrast trials resulting in attractive estimation biases towards the most frequent directions. In addition, prior expectations lead to reduced estimation variability and reaction times, as well as increased detection performance for the most frequently presented directions. When no stimulus is presented, the acquired expectations sometimes lead to false alarms (‘hallucinations’), again, mostly in the most frequent directions. Importantly, such biases were well described using a Bayesian model, where participants acquired a perceptual prior for the visual stimulus that is combined with sensory information and influences their perception. As such, this paradigm is well suited to quantitatively model variations in likelihoods and priors in individuals with ASD or schizotypal traits.

**Figure 2:**
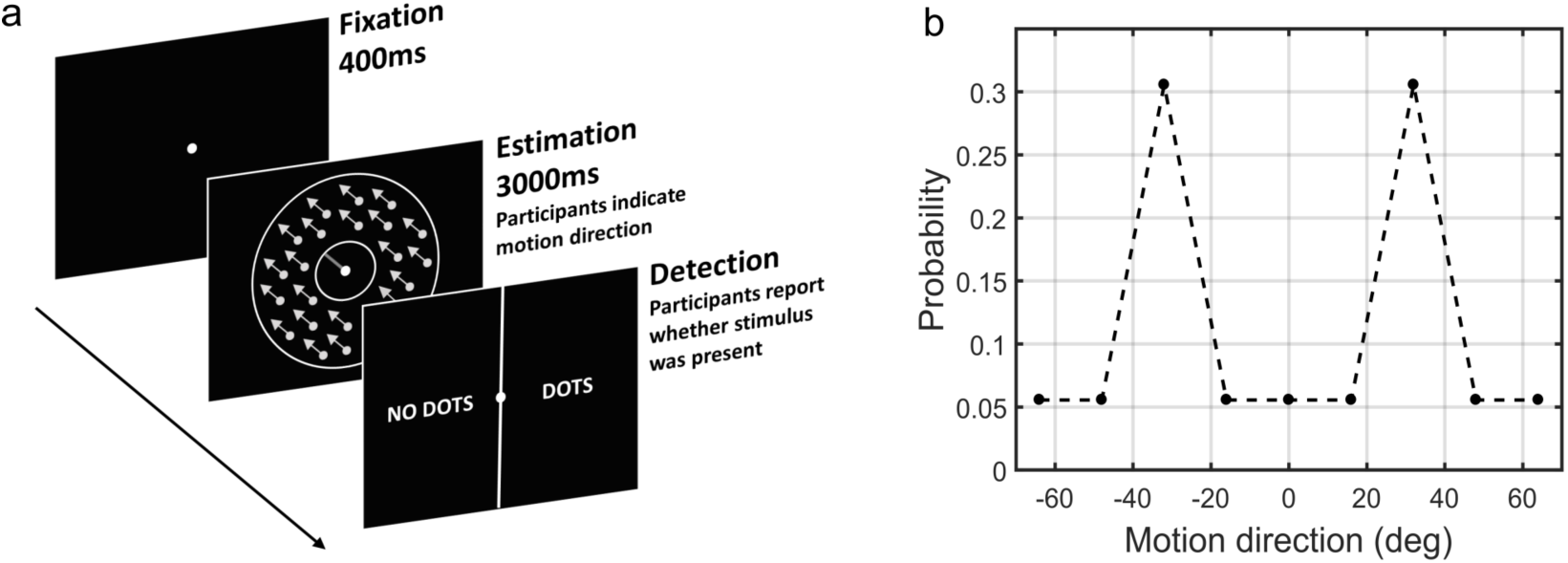
The moving dots task. (a) Sequence of events on a single trial. First, a fixation point is presented. Next, a field of coherently moving dots is presented along with an estimation bar (extending from the fixation point) which participants are required to move to indicate perceived motion direction. Lastly, in a two-alternative forced choice, participants are asked to report whether they saw the dots during the estimation part (detection task). (b) The probability of different motion directions being presented: directions at ±32° are presented more often than other directions. Motion direction is plotted relative to a central reference angle (at 0°), which was randomly set for each participant.

## Results

Here, we investigated individual differences in statistical learning in relation to autistic and schizotypal traits in a sample of 91 healthy participants. 8 participants failed to perform the task satisfactorily and were excluded from the analysis (see *Methods*), leaving 83 participants in the study (41 women and 42 men, age range: 18-69; mean: 25.7).

### Task behavior at low contrast

First, we investigated whether participants acquired priors on the group level. We discarded the first 170 trials as that is how long it took for the 2/1 and 4/1 staircases contrast levels to converge (**Supplementary Fig. 2**) and for prior effects to become significant (Section 3 in Supplementary Material). We analyzed task performance at low contrast levels (converged 2/1 and 4/1 staircases contrast levels) where sensory uncertainty is high. Replicating findings of Chalk et al. (2010), we found that on the group level people acquired priors that approximated the statistics of the task. Such priors were indicated by: attractive biases towards ±32°(**fig. 3a**) less variability in estimations at ±32° (**Fig. 3b**; standard deviation of estimations 11.9± 0.30° at ±32° versus 13.84±2.38° over all other motion directions; signed rank test: p<0.001), shorter estimation reaction times at ±32°as compared to all other motion directions (**Fig. 3c**; average reaction time was 201.87 ± 2.47 ms at ±32° versus 207.75 ± 2.60 ms over all other motion directions; signed rank test: p < 0.001) and better detection at ±32° as compared to all other motion directions (**Fig. 3d**; detected 75.57 ± 0.65% at ±32°versus 66.70 ± 0.83% over all other motion directions; signed rank test: p < 0.001).

**Figure 3:**
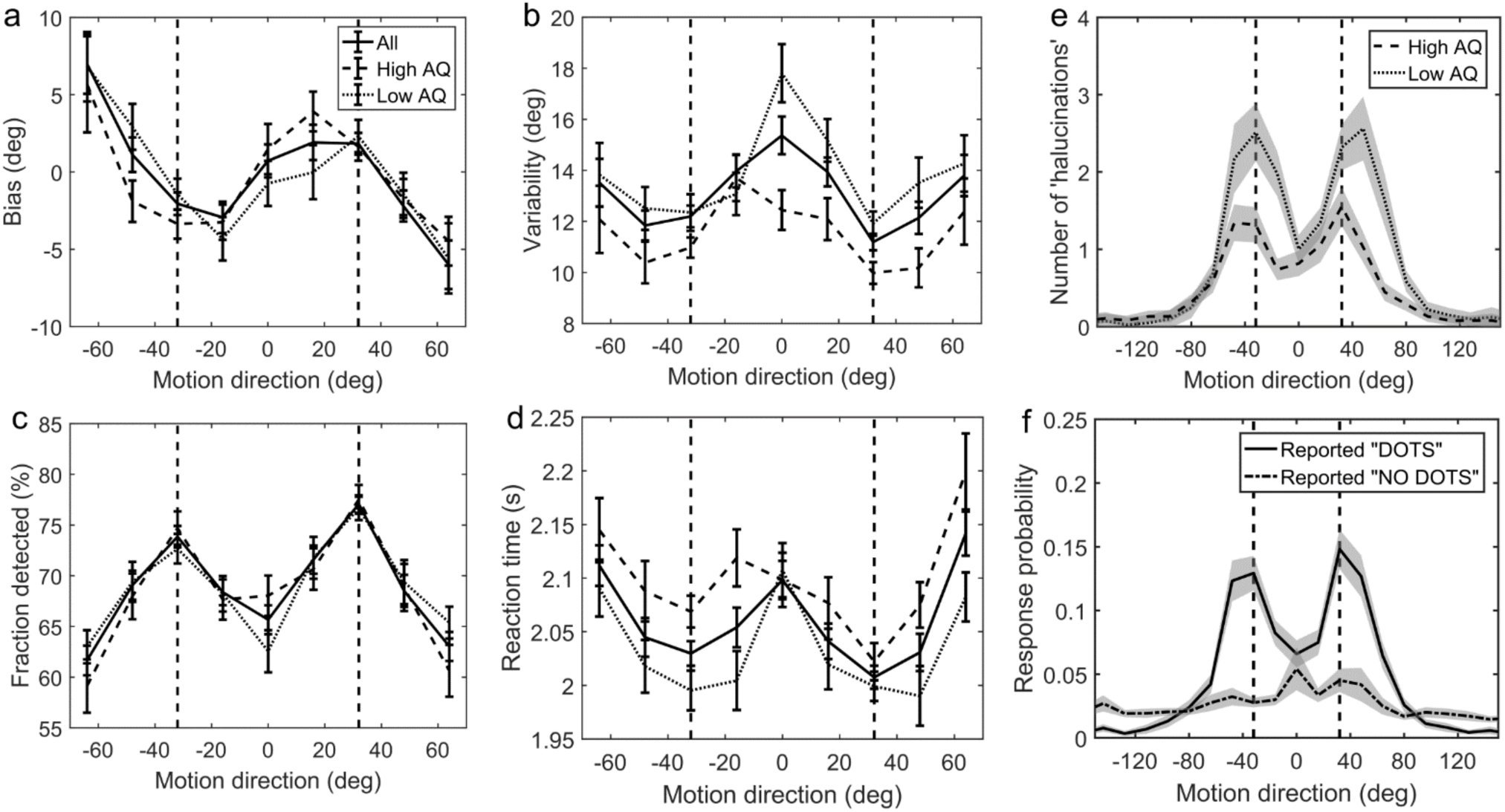
Average group performance on low-contrast trials (a-d) and on trials with no stimulus (e). (a) Mean estimation bias, (b) standard deviation of estimations, (c) estimation reaction time and (d) fraction of trials in which the stimulus was detected. (f) Probability distribution of estimation responses on trials without stimulus. The solid line denotes the estimation responses when participants reported detecting a stimulus (‘hallucinations’). The dash-dot line denotes estimation distributions when participants correctly reported not detecting a stimulus. (e) Distribution of ‘hallucinations’ for high and low AQ groups (median split). The vertical dashed lines correspond to the two most frequently presented motion directions (±32°). Error bars and shaded areas represent within-subject standard error.

### No-stimulus performance

Another indicator of acquired priors is the distribution of estimation responses on trials when no actual stimulus was presented. We found that participants sometimes still reported seeing dots (experienced ‘hallucinations’) but mostly so around ±32° (**Fig. 3f**, solid line). To quantify the statistical significance of ‘hallucinations’ around ±32°’ the space of possible motion directions was divided into 45 bins of 16° and the probability of estimation within 8° of ±32° was multiplied by the total number of bins:

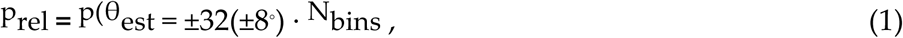

where ^N^_bins_ is the number of bins (45), each of size 16°. This probability ratio would be equal to 1 if participants were equally likely to estimate within 8° of ±32°, as they were to estimate within other bins. We found that the median of p_rel_ was significantly greater than 1 (median(p_rel_) = 1.6, p<0.001, signed rank test). Furthermore, the estimation distribution when no dots where detected (**Fig. 3f**, dash-dot line) was found to be significantly flatter (median(p_rel_) = 0, p < 0.001, signed rank test comparing with the median of p_rel_ for ‘hallucinations’), suggesting that the ‘hallucinations’ were indeed of perceptual nature (rather than related to a response bias).

### Task performance and autistic/schizotypy traits

Participants were prescreened to make sure they covered a wide range of autistic and schizotypy scores. The AQ scores in our sample ranged from 6 to 41 with a mean (±SD) of 20.3 (±8.3). The RISC scores ranged from 8 to 55 with a mean of 31.7 (±11.9), and the SPQ scores ranged from 4 to 59 with a mean of 26.4 (±13.8).

We found significant effects of autistic traits on the performance at low contrast trials: autistic traits were associated with less bias (**Fig. 4a**; mean absolute estimation bias: *ρ* = -0.228, p = 0.039) and less variability in estimations (**Fig. 4b**; mean standard deviation of estimations: *ρ* = -0.357, *p* = 0.001). In the Bayesian framework, less bias could arise either due to wider priors or narrower sensory likelihoods, while less variability could be a result of either narrower priors or narrower likelihoods (see **Fig. 1**). Thus, observing less bias and less variability together suggests that the effects are driven by narrower likelihoods. An alternative is that the differences in variability could be due to differences in motor noise, which we further assess via modeling (below). Schizotypy traits (RISC and SPQ scores) were found to have no effect on task performance at low contrast as indicated by the absence of correlations with mean absolute estimation bias (RISC: *ρ* = 0.132, p = 0.235; SPQ (N=39): *ρ* = -0.171, p = 0.298) and with mean estimation variability (RISC: *ρ* = 0.209, p = 0.058; SPQ (N=39): *ρ* = -0.253, p = 0.120); see **Supplementary Fig. 3**.

**Figure 4:**
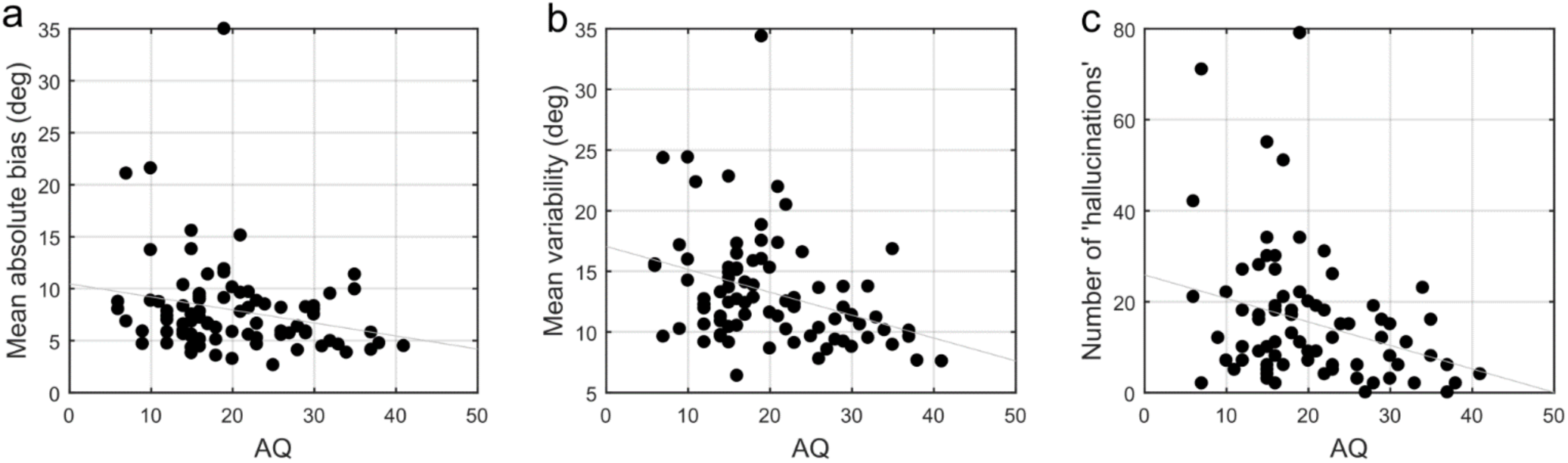
Correlations between AQ scores and task performance on low contrast trials (a, b) and when no stimulus is presented (c). (a) Mean absolute bias (ρ= -0.228, p= 0.039), (b) mean standard deviation of estimations (*ρ* = -0.357, p = 0.001), and (c) the total number of ‘hallucinations’ (*ρ* = -0.270, p = 0.014).

### No-stimulus trials and autistic/schizotypal traits

We also investigated how the traits affected performance on trials when no actual stimulus was presented. First, we looked at the total number of estimations. We found that autistic traits were associated with less ‘hallucinations’ (**Fig. 4c**; *ρ* = -0.270, p = 0.014), while schizotypal traits were found to have no effect on the number of ‘hallucinations’ (RISC: *ρ* = 0.151, p = 0.173; SPQ (N=39): *ρ* = 0.006, p = 0.971). Secondly, we looked for relationships between the traits and how the estimations on no-stimulus trials were distributed. Specifically, we were interested in whether the traits predicted how densely ‘hallucinations’ were distributed around ±32°, as this could be considered to reflect the differences in the width of the underlying acquired prior distribution. To determine this, we looked at the fraction of total ‘hallucinations’ in the region around ±32° for three different-sized windows: 1) Within 8°, within 16° and within 24° of ±32°. These measures suggested that none of the traits had any effect on how ‘hallucinations’ were distributed, suggesting no differences in the acquired prior distributions (fraction of hallucinations within 8° of ±32°: AQ - *ρ* = 0.070, p = 0.527; RISC - *ρ* = -0.106, p = 0.341; SPQ - *ρ* = 0.024, p = 0.882; within 16° of ±32°: AQ - *ρ* = - 0.093, p = 0.406; RISC - *ρ* = -0.197, p = 0.075; SPQ - *ρ* = 0.034, p = 0.837; within 24° of ±32°: AQ - *ρ* = 0.003, p = 0.977; RISC - *ρ* = -0.070, p = 0.527; SPQ - *ρ* = 0.147, p = 0.371).

### Modeling results

#### Group level results

To quantitatively evaluate the relationships between underlying perceptual mechanisms and task performance we fitted a range of generative models. One class of models was Bayesian - it was based on the assumption that participants combine prior expectations with uncertain sensory information on a single trial basis (**Fig. 5**).

**Figure 5.**
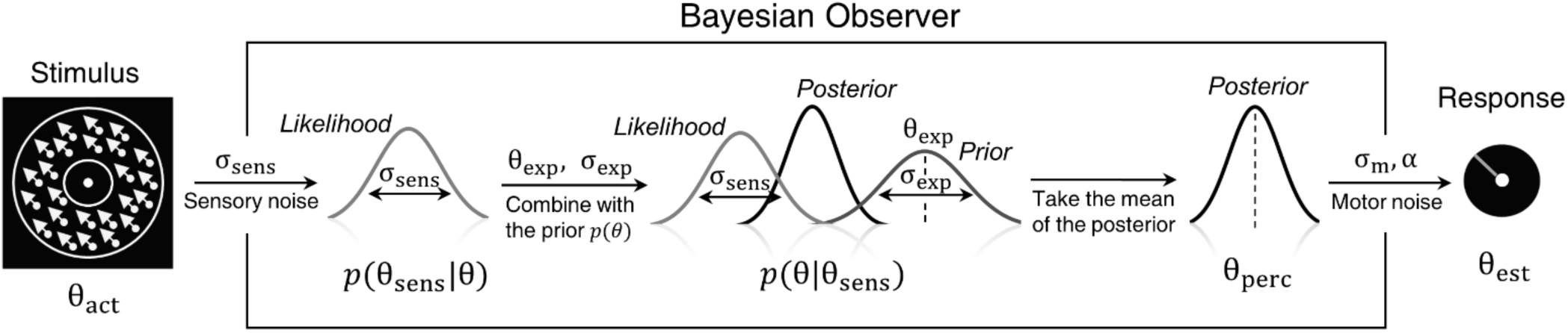
Bayesian model of estimation response for a single trial. The actual motion direction (θ_act_) is corrupted by sensory noise (σ_sens_), and then combined with prior expectations (mean θ_exp_ and uncertainty σ_exp_) to form a posterior distribution. The perceptual estimate (θ_perc_) is defined as the mean of the posterior distribution. Finally, motor noise (σ_m_) and a probability of random response () are incorporated to generate the response (θ_est_). This results in 4 free model parameters: σ_sens_, σ_exp_, θ_exp_ and *α*. Motor noise (σ_m_) is estimated from high contrast trials and is used as a fixed parameter.

To account for the possibility that the bimodal probability distribution of the stimuli, in addition to inducing prior expectations, has also affected the sensory likelihood, we constructed three variations of the Bayesian model: ‘BAYES’, where the sensory precision was constrained to be the same across all presented motion directions, ‘BAYES_varmin’, where the sensory precision was allowed to be different for the most frequently presented motion directions, but was the same across all other directions, and ‘BAYES_var’, where sensory precision was allowed to be different across all motion directions. Another class of models was based on the assumption that task performance can be explained by response strategies that do not involve Bayesian inference. That is, on any given trial participants responded based on the prior expectations or sensory information alone. We considered four variations of response strategy models: ‘ADD1’, ‘ADD2’, ‘ADD1_m’ and ‘ADD2_m’ (see Methods for details).

To compare the models, we computed BIC values for each individual for each model; we used individual BIC values as a summary statistic and compared the models using signed rank test in order to preserve individual variability (**Fig. 6a**). We found that the BAYES model had significantly smaller BIC values than the remaining models (see the p-values within **Fig. 6a**).

**Figure 6:**
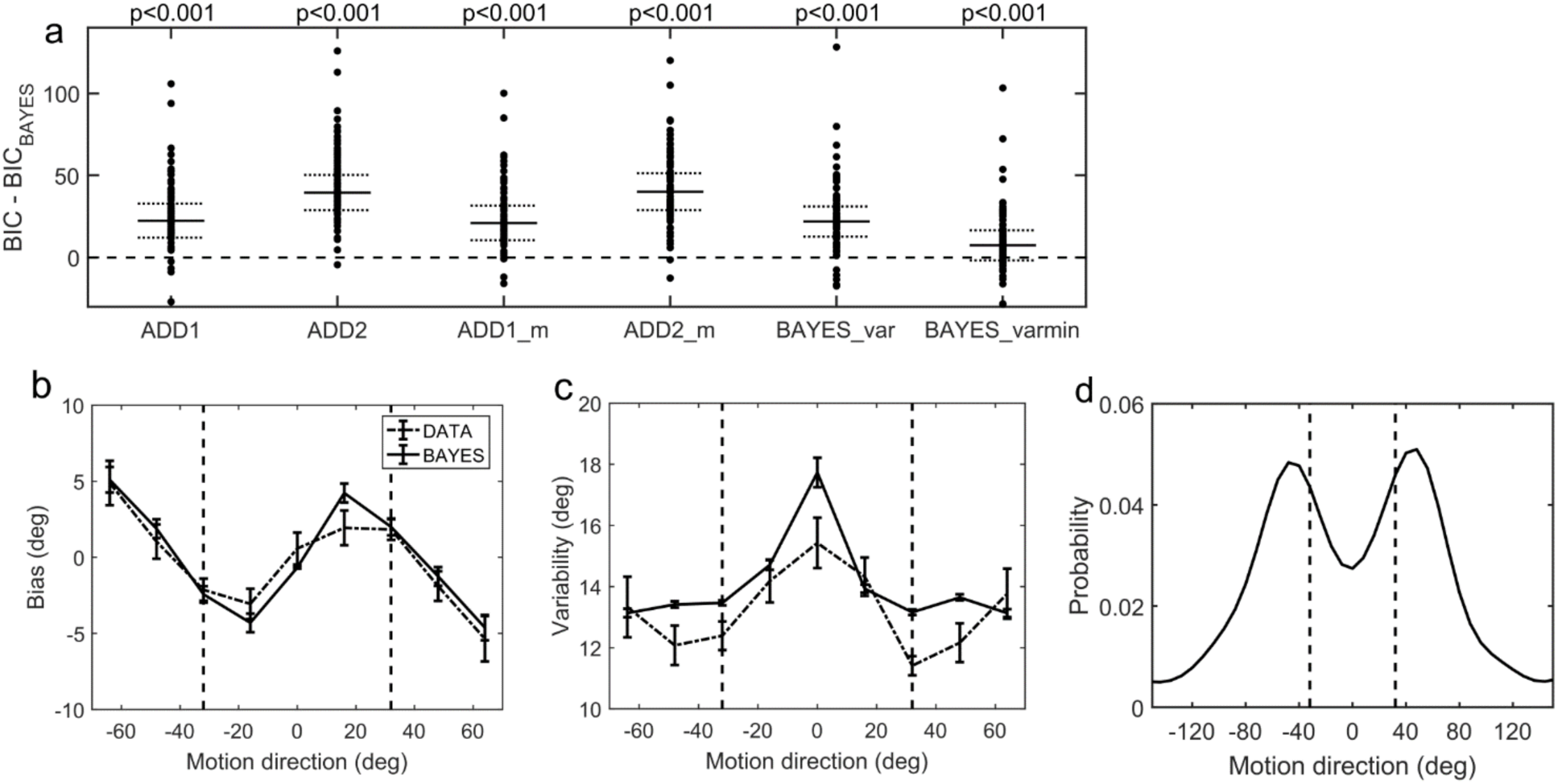
Modelling results. (a) Model comparison for all participants using Bayesian Information Criterion (BIC). y-axis measures the relative difference between BIC of each model (as indicated on the x-axis) and BIC of BAYES model. Values greater than zero on the y-axis indicate that the BAYES model provided a better fit. Each dot represents a participant. Solid horizontal lines denote median values; doted horizontal lines denote 25th and 75th percentiles. p-values above the plot indicate whether the median of the difference was significantly different from zero for each model (signed rank test). Panels (a) and (c) present task performance at different motion directions as predicted by BAYES model: (b) estimation bias, (c) standard deviation of estimations. Error bars represent within-subject standard error. (d) Population averaged prior as recovered via BAYES model. The vertical dashed lines correspond to the two most frequently presented motion directions (±32).

### Model parameters and autistic/schizotypal traits

Correlational analysis of BAYES model parameters and AQ revealed that autistic traits did not have any effect on acquiring prior expectations. There was no correlation between AQ and the mean of the acquired prior distribution θ_exp_, or between the AQ and the precision of the prior σ_exp_ (**Fig. 7a,b**; *ρ* = 0.016, p = 0.884 and *ρ* = 0.002, p = 0.987, respectively).

**Figure 7:**
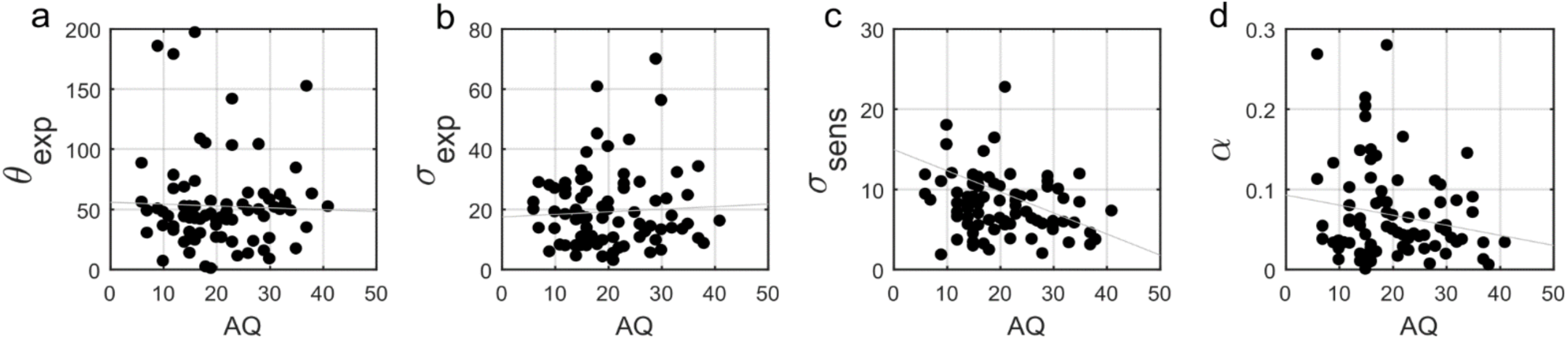
Correlations between AQ scores and BAYES model parameters. (a) θ_exp_ - mean of the prior expectations (*ρ* = 0.016, p = 0.884), (b) σ_exp_ - uncertainty of the prior distribution (*ρ* = 0.002, p = 0.987), (c) σ_sens_- uncertainty in the sensory likelihood (*ρ*=–0.285, p=0.009) and (d) *α* - fraction of random estimations (*ρ* = -0.074, p = 0.509).

Importantly, autistic traits were found to be strongly associated with less uncertainty in the sensory likelihood, σ_sens_ (**Fig. 7c**; *ρ* = –0.285, p = 0.009). Moreover, the ratio between the uncertainties of likelihood and prior was marginally significantly correlated with AQ (*ρ* = –0.211, p = 0.055), consistent with smaller estimation biases found along the autistic traits. Finally, there was no correlation with the amount of random estimations (**Fig. 7d**; *ρ* = –0.074, p = 0.509). Motor noise, which was estimated from high contrast trials, separately from all other parameters (see Methods), was also negatively correlated with autistic traits (ρ = –0.242, p = 0.028). On the other hand, consistent with the absence of differences in the behavioral findings, schizotypal traits were not associated with any difference in the BAYES model parameter values (**Supplementary Fig. 5**).

### Parameter recovery for BAYES

Finally, to further investigate that in our experimental paradigm the influence of stronger likelihoods can be distinguished from that of weaker priors ^10, 11^ we performed parameter recovery for the winning BAYES model. Parameter recovery involves generating synthetic data with different sets of parameters (‘actual parameters’) and then fitting the same model to estimate the parameters (‘recovered parameters’) that are most likely to have produced the data. If actual and recovered parameters are in a good agreement, it means that the effects of different parameters can be reliably distinguished. At the same time, parameter recovery is also affected by the parameter estimation methods and even more so by the amount of data used for model fitting. Therefore, parameter recovery provides an overall check for the reliability of modelling results and is recommended as an essential step in computational modelling approaches ^30^.

We found that overall BAYES model (and MLE parameter estimation using simplex optimization function) recovered parameters well (**Fig. 8**).

**Figure 8:**
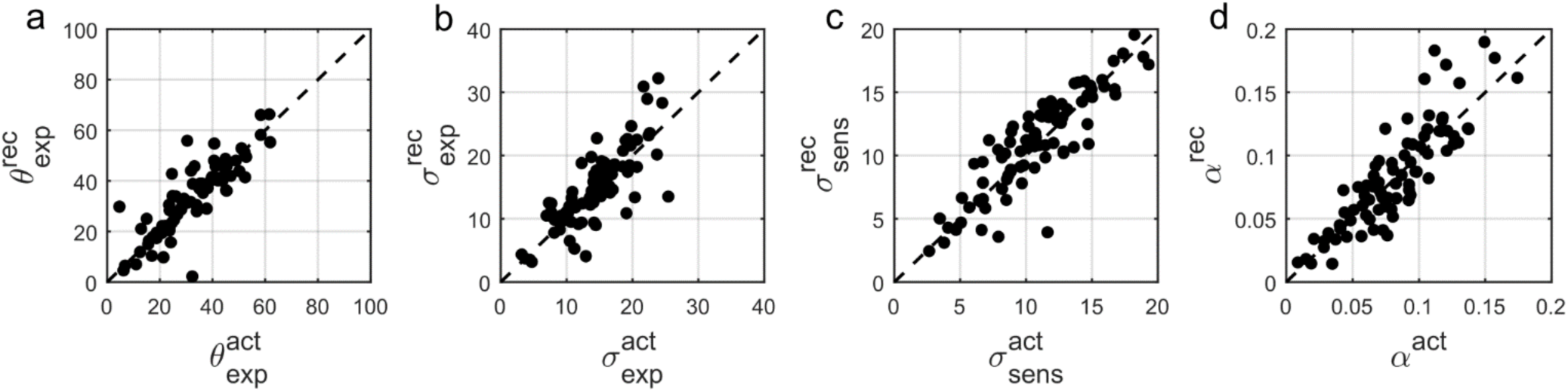
Comparison of actual (x-axis) vs. recovered (y-axis) parameters using the ‘BAYES’ model. (a) θ_exp_ - mean of the prior expectations (r = 0.77), (b) σ_exp_ - uncertainty of the prior distribution (r = 0. 68), (c) σ_sens_ - uncertainty in the sensory likelihood (r = 0.88), (d) *α* - fraction of random estimations (r = 0.87). The dashed diagonal line is a reference line indicating perfect parameter recovery.

Parameter of the highest relevance for our results, the uncertainty in the sensory likelihood, σ_sens_, was recovered most reliably (r = 0.88), followed by the fraction of random estimations, *α* (r = 0.87), the mean of the prior expectations distribution, θ_exp_ (r = 0.77) and the uncertainty in prior expectations, σ_exp_ (r = 0.68).

## Discussion

In this study, we investigated whether autistic and schizotypal traits are associated with differences in the implicit Bayesian inference performed by the brain. Specifically, we wanted to know whether autistic and schizotypal traits are accompanied by 1) differences in how the priors are updated and/or in their precision and/or by 2) differences in the precision with which the sensory information (the likelihood) is represented. We used a visual motion estimation task ^29^ that induces implicit prior expectations via more frequent exposure of two motion directions (±32°). We found that on the group level (N=83) participants acquired prior expectations towards ±32° motion directions. This was indicated by shorter estimation reaction times and better detection at ±32°, as well as attractive biases towards ±32° and reduced estimation variability at ±32°. Moreover, when no stimulus was presented, participants sometimes still reported seeing the stimulus, mostly around ±32°. Performance was best explained by a simple Bayesian model, which provided a good fit to the data and captured the characteristic features of perceptual bias and variability. This model provided estimates of Bayesian priors and sensory likelihoods for each participant, which were then analyzed in relation to participants’ schizotypal and autistic traits.

Schizotypal traits were found to have no measurable effect on perceptual biases in our task and, therefore, were not associated with any differences in the precision ascribed to priors and likelihoods. This finding challenges recent accounts of positive symptoms of schizophrenia that predict impaired updating of priors and an imbalance in precision ascribed to sensory information and prior expectations ^1–3^. An immediate explanation might be that the influence of schizotypal traits in the healthy population is not strong enough to lead to behavioral differences, even if the dimensionality assumption holds. It is likely for example, that, even if they sometimes scored high in schizotypal traits, our participants didn’t experience daily hallucinations. That they would not exhibit an overweighting of perceptual priors would then be consistent with the recent study of Powers et al^31^. However, our results also contradict recent findings by Teufel et al.^16^ who found that both early psychosis and schizotypal traits are associated with a relatively stronger influence of prior knowledge. In that study, participants were presented with ambiguous two-tone versions of images before and after seeing the actual images in full color and had to report whether the presented two-tone image contains a face. People with stronger schizotypal traits and early psychosis showed a larger improvement after seeing the color images. A possible difference between their study and ours might be the level at which the priors operate: the low-level prior for basic perceptual features (as induced in our task) might function at a hierarchically lower level than prior knowledge related to complex collection of features and semantic content (faces). The level at which prior expectations are induced has indeed been shown to matter. A series of studies by Schmack et al^17, 27, 28^ using 3D rotating cylinders report weaker low-level (perceptually-induced - stabilizing) priors but stronger high-level (cognitively-induced) priors in both schizophrenia and schizotypal traits. It is difficult to compare and reconcile these findings with ours. One possibility is that the priors induced in our task lie in between their perceptual and cognitive levels. The taxonomy of priors in relation to their place in the computational hierarchy or to their complexity or specificity is still far from being established ^32^ and thus the potential relevance of such distinctions is still not known. Autistic traits were associated with significant behavioral differences: weaker biases and lower variability of direction estimation on low contrast trials. Modeling revealed that this was because of increased sensory precision as well as a reduction in motor noise, while there was no attenuation of acquired priors. Parameter recovery analysis confirmed that our methodology provides reliable parameter estimates and, in particular, allows disentangling variations in priors and likelihoods.

Autistic traits were also found to be associated with less false detections (‘hallucinations’) on trials when no stimulus was presented, consistent with the idea that prior expectations had less influence in individuals with higher AQ. In an attempt to measure those individual differences, we fitted a more sophisticated Bayesian model that could account not only for the estimation performance but also for the detection data (see **S4** in Supplementary Material). This model provided a good fit to both estimation and detection data, and preserved the correlation between ASD traits and the precision of the motion direction likelihood (ρ= -0.235, p= 0.032). However, parameter recovery was not as good as for the BAYES model presented above (see **Supplementary Fig. 11**) and for this reason we focused on the simpler model in this paper.

Overall, our findings are in agreement with most of the recent Bayesian theories of ASD, namely, that autistic traits are associated with a relatively weaker influence of prior expectations. However, we find that this is due to enhanced sensory precision^6, 7, 10, 13^, rather than attenuated priors per se^4^. Other empirical studies inspired by the Bayesian accounts have reported either attenuated or intact priors, but most are subject to methodological limitations, either because they did not use computational modeling^15, 22–24^ or because their model could not extract likelihoods and quantify their variations^14, 26^.

The idea that sensory processing could be enhanced in autism has long been proposed outside the Bayesian framework. Autistic traits have been associated with enhanced orientation discrimination^33^, but only for first-order (luminance-defined) stimulus^34^. This enhancement has been proposed to be a result of either enhanced lateral ^34^, or a failure to attenuate sensory signals via top-down gain control^6^, both of which could be directly related to narrower likelihoods in the Bayesian framework^35^. However, in motion perception, previous research did not find improved discrimination for first-order stimulus in autism, while for second-order (texture-defined) stimulus, the autistic group was found to underperform^36^. Our findings challenge these results and call for more research in this area.

In ASD as in schizotypy, prior integration might function differently at different levels of sensory processing. For example, Pell et al.^23^ reported intact direction-of-gaze priors for healthy individuals with high autistic traits and for highly functional individuals with a clinical diagnosis. The authors did not directly investigate differences in sensory precision, but the lack of behavioral differences suggests that there was none. Arguably, their paradigm involves more complex stimuli than used in our task, which are also strongly associated with semantic content (faces). It would not be surprising if increased sensory precision does not extend to such stimuli. In fact, autistic individuals are known to exhibit differential performance based on the complexity of the stimulus^34^, which also lies at the foundation of some theoretical accounts, such as the ‘Weak Central Coherence’^37^.

In our paradigm people acquire prior expectations very quickly, within 200 trials (see Section 3 in Supplementary material), which did not allow us to study individual differences in the rate at which the priors are acquired. Bayesian accounts predict differences in the dynamical updating of the priors, namely, that both autistic and schizotypal traits should be associated with increased learning rate - which is the ratio of likelihood and posterior precisions ^7^. Our findings of increased sensory precision in autistic traits also suggest that their learning rate should be faster. Future work will aim at directly testing this.

Another aspect that our paradigm could not test is the specificity of the acquired priors ^32^. Some Bayesian accounts ^5^ predict that priors may be overly context-sensitive in autism. This is in line with the view that generalization is impaired in autism ^38^. Furthermore, such over-specificity is thought to be stronger with more repetitive stimuli ^39^. Future research could address this using statistical learning paradigms that incorporate increasingly distinct contexts or stimuli.

## Conclusion

We investigated statistical learning and Bayesian inference in a visual motion perception task along autistic and schizotypal traits. To our knowledge, this study is the first to investigate differences in Bayesian inference along both trait spectra in a single task. Furthermore, this study is the first visual study to computationally disentangle and quantitatively assess the variations in individuals’ likelihoods and priors. Surprisingly, schizotypal traits were found to have no effect on task performance and thus were not associated with any differences in the underlying statistical learning and Bayesian inference. For autistic traits, however, significant behavioral differences in prior integration were found, which were due to an increase in the precision of internal sensory representations in participants with higher AQ.

## Methods

### Participants

91 (47 females, 44 males, age range: 18-69) naïve participants with no motor disabilities and with normal (or corrected to normal) vision were recruited from the general population. We advertised for participants using posters and the internet across University of Edinburgh locations and other sites across Edinburgh. All participants gave informed written consent and received monetary compensation for participation. The study was approved by the University of Edinburgh School of Informatics Ethics Panel.

### Questionnaires

ASD was assessed using 50-item version Autism Spectrum Quotient (AQ) ^40^, which is commonly used for assessing milder variants of autistic-like traits within the general population. Schizotypal traits were assessed using The Rust Inventory of Schizotypal Cognitions (RISC) ^41^. RISC is specifically developed to measure schizotypal traits in the general population. In addition, a sub-group of 41 participants also completed Schizotypal Personality Questionnaire (SPQ) ^42^. Finally, all participants were also asked to complete the Warwick-Edinburgh Mental Well-being Scale (WEMWBS)^43^ in order to control for potential depression-induced differences in performance ^44^.

### Apparatus

The visual stimuli were generated using Matlab Psychophysics Toolbox ^45^. Participants viewed the display in a dark room at a distance of 80-100cm. The stimuli consisted of a cloud of dots with a density of 2 dots/deg^2^ moving coherently (100%) at a speed of 9°/sec. Dots appeared within a circular annulus with minimum diameter of 2.2° and maximum diameter of 7°. The stimuli were displayed on a Dell P790 monitor running at 1024x768 at 100 Hz. The display luminance was calibrated using a Cambridge Research Systems Colorimeter (ColorCal MKII).

### The task

The task was developed previously in our laboratory^29^. Participants have to: i) estimate the direction of coherently moving simple stimuli (dots) that are presented at low contrast levels (estimation task) and then ii) indicate whether they have actually perceived the stimulus or not (detection task). Since Chalk et al.^29^ had shown that the effects of acquired priors become significant within the first 200 trials, instead of two experimental sessions of 850 trials each as in the original study, we used a single session of 567 trials (lasting around 40 min).

Each trial started by first displaying a fixation point (0.5°, 12.2 cd/m^2^) for 400 ms, after which a field of moving dots appeared along with an orientation bar (length 1.1°, width 0.03°, luminance 4 cd/m^2^, extending from the fixation point). Initial angle of the bar was randomized for each trial. Participants had to estimate the direction of motion by aligning the bar (using a computer mouse) to the direction the dots were moving in, and by clicking the mouse button to validate their estimate. The display cleared when either the participant had clicked the mouse or when 3000 ms had elapsed. On trials where no stimulus was presented, the bar still appeared for the estimation task to be completed.

After a 200ms delay, the participants had to indicate whether they had actually detected the presence of dots in the estimation period (detection task). The display was divided into two parts by a vertical white line across the center of the screen, the left hand side area reading “NO DOTS” and the right hand side area reading “DOTS” (**Fig. 2a**). The cursor appeared in the center of the screen, and participants had to move it to the left or right and click to indicate their response. Immediate feedback for correct or incorrect detection responses was given by a cursor flashing green or red, respectively. The screen was cleared for 400 ms before the start of a new trial. Every 20 trials, participants were presented with feedback on their estimation performance in terms of average estimation error in degrees (e.g., “In the last 20 trials, your average estimation error was 23°”). Every 170 trials (i.e. on three occasions) participants were given a chance to “have a short break to rest their eyes”, in order to prevent fatigue. Participants clicked when they were ready to continue.

### Design

The stimuli were presented at four different levels of contrast: 0 contrast (no-stimulus trials), 2 low levels contrasts and high contrast, randomly mixed across trials. There were 167 trials with no stimulus. The 2 low levels of contrast were determined using 4/1 and 2/1 staircases on detection performance^46^. There were 243 trials following the 4/1 staircase and 90 trials following the 2/1 staircase. The remaining 67 trials were at high contrast, which was set to 3.51 cd/m^2^ above the background luminance.

For the two low contrast levels, there was a predetermined number of possible directions: 0°, ±16°, ±32°, ±48°, and ±64° with respect to a reference direction. The reference direction was randomized for each participant. For the 2/1 staircased contrasts, each predetermined motion direction was presented equally frequently. Unbeknownst to participants, stimuli at high and 4/1 staircase contrasts were presented more frequently at -32° and +32° motion directions, resulting in a bimodal probability distribution (**Fig. 1b**). For the 4/1 staircase contrast level, the dots were moving at ±32° in 173 (~70%) trials and in all the other predetermined motion directions in the remaining 70 (~30%) trials equally frequently. At the highest contrast level, 34 (~50%) trials had the dots moving at ±32° and the remaining 33 (~50%) trials were at random directions (i.e. not just the predetermined directions).

### Data analysis

Responses on high contrast trials were used as a performance benchmark to ensure that participants were performing the task adequately. 8 out of 91 participants failed to satisfy pre-defined performance criteria (at least 80% detection and less than 30° root mean squared error of estimations) and were excluded from further analysis (**Supplementary fig. 1**).

Data analysis on the estimation of motion directions was performed on 4/1 and 2/1 staircased contrast levels only and only on trials where participants both validated their choice with a click within 3000 ms in the estimation part and clicked “DOTS” in the detection part. The first 170 trials of each session were excluded from the analysis, as this was the upper limit for the convergence of the staircases to stable contrast levels (**Supplementary Fig. 2**).

After removing these trials, the luminance levels achieved by the 2/1 and 4/1 staircases were found to be considerably overlapping (**Supplementary Fig. 2**). Therefore, the data for both of these contrast levels was combined for all further analysis.

To account for random estimations (either accidental or intentional) that participants made on some trials, we fitted each participant’s estimation responses to the probability distribution:

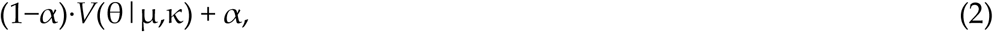

Where *α* is the proportion of trials in which participant makes random estimates, and *V*(θ|μ,κ) is the probability density function for the estimated angle *θ* for von Mises (circular normal) distribution with the mean *μ* and variance 1/*κ*. The parameters *μ* and *κ* of the von Mises distribution were determined by maximizing the likelihood of the distribution in Eq. (2) for each presented angle.

To analyze the distribution of estimations in no-stimulus trials, we constructed histograms of 16° size bins. These histograms were converted into probability distributions by normalizing over all motion directions. We analyzed the estimation distribution when participants reported seeing dots (clicked “DOTS”) within no-stimulus trials. We interpreted these false alarms as a simple form of perceptual “hallucination”.

### Modelling

#### Bayesian models

Bayesian models assume that participants combined a learned prior of the stimulus directions with their sensory evidence in a probabilistic manner. We first assume that participants make noisy sensory observations of the actual stimulus motion direction (θ_act_), with a probability

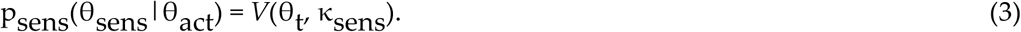

where θ_t_ itself varies from trial to trial around θ_act according to_ p(θ_t_ | θ_act_) = *V*(θ_act_, κ_sens_).

While participants cannot access the “true” prior, P_exp_(**θ**), directly, we hypothesized that they learned an approximation of this distribution, denoted P_exp_(**θ**). This distribution was parameterized as the sum of two von Mises distributions, centered on motion directions ®exp and θ_exp_, and each with variance 1/κ_exp_:

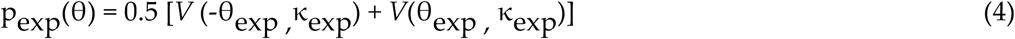

Combining these via Bayes’ rule gives a posterior probability that the stimulus is moving in a direction θ:

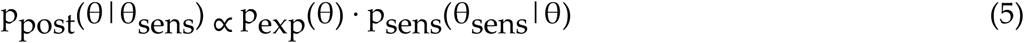

The perceived direction, θ_perc_, was taken to be the mean of the posterior distribution (almost identical results would be obtained by using the maximum instead). Finally, we accounted for motor noise and a possibility of random estimates on some trials via:

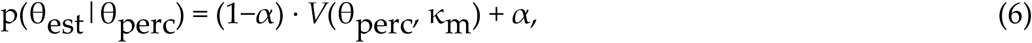

where *α* is the proportion of trials in which participants make random estimates and 1/κ_m_ is the variance associated with motor noise.

Increased exposure to some motion directions might not only give rise to prior expectations, but also affect the likelihood function ^47^. Therefore, we fitted two more model variants: ‘BAYES_var’ where κ_sens_ varied with the stimulus direction (i.e. it took five different values for each of the angles: 0_°_, ±16_°_, ±32_°_, ±48, ±64) and ‘BAYES_varmin’ where κ_sens_ was allowed to be different for ±32_°_ but was the same for all other directions.

### Response strategy models

We wanted to test whether task behavior might be better explained by simple behavioral strategies. This class of models assumed that on trials when participants were unsure about the presented motion direction, they made an estimation based solely on prior expectations, while on the remaining fraction of trials they made unbiased estimates based solely on sensory inputs. The first model, ‘ADD1’, assumed that estimations derived from prior expectations were simply sampled from a learntexpected distribution, pexp(θ)(see Chalk et al.^29^ and **Supplementary Information**). The second model, ‘ADD2’, was just as ‘ADD1’ except when participants were unsure about the stimulus motion direction, instead of sampling from the complete learned probability distribution ranging from -180° to +180°, they effectively truncated this distribution on a trial by trial basis and sampled from only one part of it, negative (-180° to 0°) or positive (0° to +180°), depending on which side of the distribution the actual stimulus occurred (see Chalk et al, 2010 and SI). We also considered slight variations of the ‘ADD1’ and ‘ADD2’ models, denoted ‘ADD1_m’ and ‘ADD2 m’ respectively. These were identical to ‘ADD1’ and’ADD2’ except from setting 1/κ_exp_ to zero; that is, on trials when perceptual estimates were derived only from expectations, they were equal to the mode of the learnt distribution (i.e. no uncertainty).

### Parameter estimation

We used performance in high contrast trials to estimate motor noise, 1/κ_m_, for each individual. We assumed that, for those trials, sensory uncertainty was close to zero (1/κ_sens_ ≈ 0). Motor noise was then determined by fitting estimation responses to the distribution in Eq. (2) by replacing μ with the actual motion direction, θ_act_ The estimated motor noise was used in all subsequent model fitting as a fixed parameter. The rest of the free parameters were estimated by fitting the response data at the two low (staircased) contrast levels. For each model with a set of free parameters *M*, we computed the probability distribution p(θ_est_|θ_act_; M) of making an estimate θ_est_ given the actual stimulus direction θ_act_. For the response strategy models, by definition, the p(θ_est_|θ_act_; M) corresponds to average behavior in the task.

The parameters were estimated by maximizing the fit of the log likelihood function for the experimental data for each participant individually. The maximum likelihood was found using a simplex algorithm, using “fminsearchbnd” Matlab function. To avoid convergence at a local maximum we constructed a grid of initial κ_exp_ and κ_sens_ parameter values covering the range found in previous studies. We selected the resulting set of parameters that corresponded to the largest log-likelihood.

### Model Comparison

To compare the model fits we used Bayesian Information Criterion (BIC), which approximates the log of model evidence ^48^:

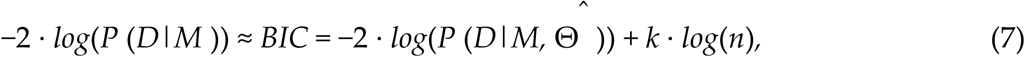

where M is model, D is observed data and *P* (D|*M*, Θ^^^) is the likelihood of generating the experimental data given the most likely set of parameters, Θ^^^; *k* is the number of model parameters and *n* is the number of data points (or equivalently, the number of trials). BIC evaluates the model by how it fits the data by also penalizing for model complexity (number of parameters); lower BIC score indicates a better model.

### Parameter recovery

To determine whether the BAYES model can distinguish the effects of strong likelihoods from those of weak priors ^10, 11^ and to evaluate the robustness of our methods, we performed parameter recovery. First, we generated 80 sets of parameters (i.e. 80 synthetic individuals) by randomly sampling each parameter from a Gaussian distribution centered on the mean value of each parameter found in our sample (40° for θ_exp_, 15° for σ_exp_, 10° for σ_sens_, 0.06 for *α* and 10^°^ for σ_m_. Second, for each set of parameters, we simulated data for 200 trials with the Bayesian model by randomly sampling from the estimation probability distribution. We used 200 simulated trials only, to match the empirical data (200 corresponds to the amount of experimental trials used for fitting, after excluding high contrast and zero contrast trials)^1^. Finally, we fitted the BAYES model to the simulated data. To evaluate the goodness of recovered parameters, we computed Pearson’s correlation between the actual parameters and the recovered parameters.

### Statistical tests

Due to the presence of outliers in our data, we used Spearman’s correlations for measuring the strength of the effects. We have also used Wilcoxon signed rank test for repeated measures analysis.

## Acknowledgements

We thank Gizem Aras for assisting in data collection, and Katie Richards for assisting with participants’ recruitment. This work was supported by NARSAD Young Investigator grant number 19271 (to PS). PS and AS were also supported by NIH grant # 1R01EY023582.

## Competing financial interests

The authors declare no competing financial interests.

## Footnotes

1 Simulating more trials would result in a better parameter recovery but the results would no longer be informative about the reliability of parameters estimated from empirical data.

